# Autophagy maintains the homeostatic environment in the male reproductive accessory organs playing a key role in fertility

**DOI:** 10.1101/2023.07.21.549845

**Authors:** Adil Jaulim, Liam D Cassidy, Andrew RJ Young, Adelyne SL Chan, Anne Y Warren, Angela E Taylor, Wiebke Arlt, Guochen Lan, Martyn L Blayney, Olivia Davidson, Christopher LR Barratt, Simon Pacey, Masashi Narita

**Affiliations:** Cancer Research UK, Cambridge Institute, University of Cambridge, Cambridge, CB2 0RE, UK; University College London Hospitals NHS Foundation Trusts, Department of Cellular Pathology, London, W1T 4EU, UK; Department of Histopathology, Cambridge University Hospitals NHS FT, Cambridge, CB2 0QQ, UK; Institute of Metabolism and Systems Research, University of Birmingham, Edgbaston, Birmingham, B15 2TT, UK; Medical Research Council Laboratory of Medical Sciences, London, W12 0NN, UK; Institute of Clinical Sciences, Imperial College London, London, W12 0NN, UK; Stem Cell and Regenerative Consortium Centre, University of Hong Kong, China; Bourn Hall Clinic, High St, Bourn, Cambridge, CB23 2TN, UK; Division of Systems Medicine, School of Medicine, Ninewells Hospital and Medical School, University of Dundee, Dundee, DD1 9SY, UK; Department of Oncology, Clinical School, University of Cambridge, Cambridge, CB2 0QQ, UK

## Abstract

Autophagy has been implicated in male fertility but its specific role in the post-testicular organs remains unclear. Here, we investigate this in mice expressing a doxycycline-inducible RNAi against Atg5 (Atg5i). Systemic autophagy inhibition in Atg5i mice resulted in the morphological and functional abrogation of the male accessory sex organs, leading to male subfertility. However, the testis was largely protected, likely due to the limited permeability of doxycycline through the blood-testis barrier. Interestingly, restoration of autophagy by doxycycline withdrawal in Atg5i mice led to substantial recovery of the phenotype in the accessory organs. This model offers a unique opportunity to dissect the pre- and post-testicular roles of autophagy, highlighting the non-autonomous impact of autophagy on male fertility.

## Introduction

Abnormal semen parameters are indicative of a role for male-factor infertility in around half of involuntarily childless couples (Jungwirth et al. 2012) Moreover, many studies have suggested a decreasing trend in sperm concentration over the last century (Skakkebæk et al. 2022; Virtanen et al. 2017; Swan et al. 2000). Although hampered by inherent variability in the data (Bonde and Velde 2017) and the difficulties in standardising the sampled populations, an emerging consensus of estimates seems to be that related parameters such as sperm count and concentration have dropped by around 1% annually in studies spanning periods over the last four decades (Levine et al. 2022; Huang et al. 2017; Tiegs et al. 2019). While this remains controversial, the trend of these estimates is nonetheless concerning and there does seem to be a consensus that sperm counts are hovering around a critical threshold, after which fertility may drop substantially (Daumler et al. 2016; Herrera et al. 2021). However, despite its widespread prevalence, male infertility is often overlooked as a clinical disorder by the general population (Daumler et al. 2016).

Research studies into male infertility are often focused on the role of spermatogenesis in the testes, however the auxiliary sex organs, such as the epididymis and seminal vesicles also play key roles in sperm maturation and nourishment. Here spermatozoa (sperms) develop properties essential for the fertilisation of an oocyte with transit time through the epididymis considered a critical factor (ORGEBIN-CRIST 1967). Additionally, many other functions are also attributed to the epididymis, such as sperm concentration (Belleannée et al. 2012), immunoprotection of the male gamete, and its role as a sperm reservoir.

Factors such as age and obesity are associated with declines in semen parameters (Santi et al. 2023; Silva and Anderson 2022), whilst advancing age is associated with loss of sperm volume, decreased sperm motility and increased DNA fragmentation (Sharma et al. 2015). Indeed, the association between paternal age at conception, and the risks of spontaneous abortion suggest that a reduction in sperm quality with age may impact more than just the ability to conceive (Nguyen et al. 2019).

Autophagy is a bulk cellular degradation pathway with direct links to both ageing and metabolism (Aman et al. 2021). The rate of autophagy is thought to decline with age (Cassidy and Narita 2022), whilst basal autophagy may have insufficient capacity to cope with increased demand during some stress conditions, such as obesity (Lim et al. 2014). Cell-type specific and constitutive knockout models of autophagy have previously shown a profound defect in spermatogenesis, leading to male infertility (Liu et al. 2016; Wang et al. 2014; Yoshii et al. 2016). However, the role of autophagy in the male accessory sex organs and its effect on fertility, separate from spermatogenesis, has not been investigated.

Here we employ an inducible model of autophagy inhibition, which retains autophagy in the seminiferous tubules, where spermatogenesis occurs, whilst simultaneously down-regulating autophagy in the sexual accessory organs. In these male mice, loss of autophagy was associated with a reduction in fertility, as determined by both natural copulation and *in vitro* fertilisation studies, that coincided with histological alterations to the epididymis and seminal vesicles. Restoration of autophagy and reversibility of the accessory sex organ alterations was associated with the rescue of fertility, suggesting that autophagy may play a critical role in fertility outside of spermatogenesis. Together this raises the possibility that autophagy promoting therapies may represent an avenue for some forms of male factor subfertility in situations wherein spermatogenesis is intact.

## Results and Discussion

### Body-wide inhibition of autophagy leads to a shrinkage of the accessory sex organs

To investigate the effects that a reduction in autophagy would have on the mature urogenital system, we used our Atg5i mouse model, which harbours a doxycycline (dox)-inducible shRNA against *Atg5*, an essential gene for autophagy (Cassidy et al. 2018, 2020). Whole-body autophagy deficiency often causes early mortality due to severe neurotoxicity (Kuma et al. 2017). However, the limited permeability of dox through the blood-brain barrier (BBB) means our Atg5i mice (Dow et al. 2012; Cassidy et al. 2020) exhibit no discernible neurotoxic effects upon systemic RNAi activation, thereby providing an opportunity to evaluate the long-term effects of systemic autophagy reduction (Cassidy et al. 2020). Sexually mature eight-week-old male Atg5i mice were placed on a dox diet for six weeks (Fig. 1A). In agreement with constitutive Atg5 KO mice that also harbor brain-specific expression of *Atg5* (Atg5^-/-^;NSE- Atg5 mice), and hence do not succumb to neonatal death (Yoshii et al. 2016), Atg5i mice displayed a significant reduction in the size and weight of the epididymis, seminal vesicles (SV), and anterior prostate (Fig. 1B). The anterior prostate (AP) appeared as a thin translucent ‘membrane-like’ structure (Supplemental Fig. S1A).

**Fig. 1.**
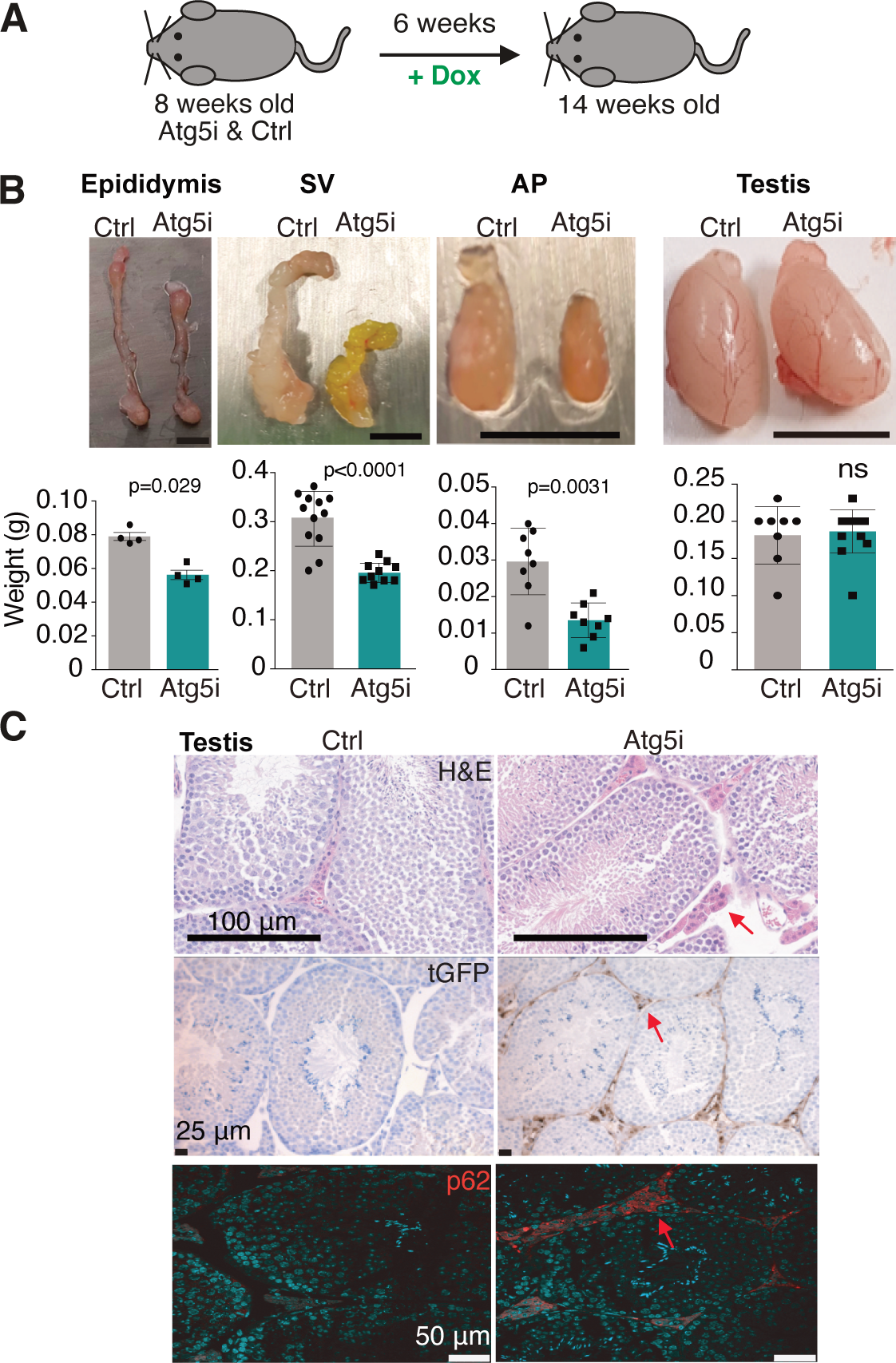
Loss of Autophagy induces shrinkage of the accessory sex organs. (*A*) Young adult male Atg5i or control mice were treated with doxycycline (dox) to induce autophagy inhibition body-wide for 6 weeks. (*B*) Loss of autophagy leads to a castration-like phenotype (SV; Seminal Vesicles, AP: Anterior Prostate). Values are mean +/- SEM. Student’s t-test. (*C*) Seminiferous tubules do not show evidence of autophagy inhibition. Representative IHC images for tGFP, which linked to sh-Atg5 expression, and p62. Only Leydig cells display evidence of tGFP and p62 accumulation (arrows).

However, and in contrast to autophagy knockout mice, we observed no alterations in testis weight between control and experimental genotypes (Fig. 1B). Furthermore, Atg5 protein levels appeared unchanged in whole tissue extracts of testes (Supplemental Fig. S1B). To investigate this further and ascertain whether the dox-inducible locus was activated, we performed immunohistochemistry for turbo-GFP (tGFP), which is linked to the expression of the Atg5-shRNA. Accordingly, we found activation appears localised to the interstitial space and absent inside the seminiferous tubules based on the staining pattern of tGFP expression staining (Fig. 1C). The autophagy adaptor protein p62 is a substate of autophagy and, thus, accumulates as a major component of ‘aggresomes’ in autophagy-deficient conditions. In agreement with the tGFP staining, p62 aggregation occurred only within the interstitial lining and Leydig cells (Fig. 1C), whilst histological analysis of the seminiferous tubules suggested no apparent difference between control and Atg5i mice, with numerous spermatids present. Combined, these data suggest that autophagy inhibition is spatially restricted in the testis. In much the same way that the BBB leads to no neurological issues in Atg5i mice, we posit that the blood-testis barrier (BTB) may limit doxycycline concentrations reaching the levels required for shRNA expression inside the seminiferous tubules. Although autophagy is essential for intact spermatogenesis (Wang et al. 2021), its role in post-testicular organs in fertility remains unclear. We hypothesised that such a spatial restriction of sh-Atg5 activity in the male reproductive organs may enable us to determine if the loss of autophagy primarily in the accessory organs may affect fertility separately from the initial stages of sperm formation (Wang et al. 2014; Yoshii et al. 2016).

### Epididymis

The epididymis is essential for post-testicular sperm maturation. Histological analysis of the caput of the epididymis showed that the ducts had undergone apparent hyperplasia in Atg5i mice: the epithelial lining of the ducts was one-cell thick in control mice but, when autophagy was inhibited, the duct lining appeared hyperplastic and pseudostratified, with the presence of more principal cells, the major cell type of the epididymis (Fig. 2A). The epididymal corpus in Atg5i mice displayed a high level of stromal fibrosis, as seen by PicroSirus Red staining (Fig. 2B). Western blotting confirmed a strong reduction in Atg5 levels as well as a reduction in the conversion of LC3-I to the membrane-bound LC3-II form, indicative of autophagy inhibition (Supplemental Fig. S1B). Interestingly, immunohistochemistry for autophagy substrates, LC3 and p62 highlighted a general build-up of p62 (Fig. 2D, Supplemental Fig. S1C), with clear cells particularly sensitive to this accumulation (Fig. 2D, Supplemental Fig. S1D). Clear cells are known to be highly enriched for genes involved in membrane trafficking and cellular metabolism (Rinaldi et al. 2020). Clear cells, together with other epithelial cells, such as principal cells, play a key role in the maintenance of the luminal environment.

**Fig. 2.**
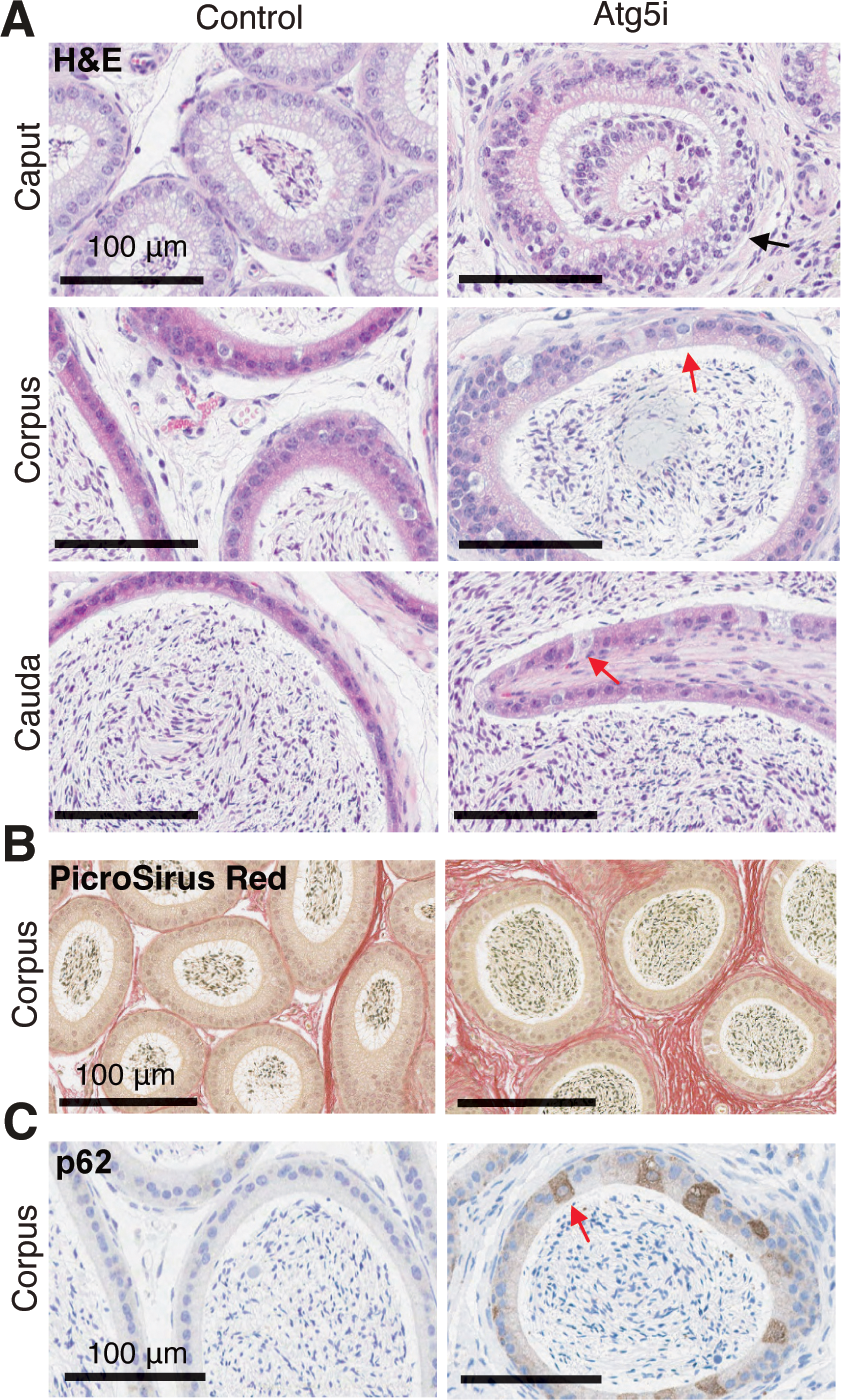
Atg5 knockdown leads to morphological changes in the epididymis. (*A*) H&E staining of indicated regions of the epididymis (6w on dox). Black arrow, increased cellularity of epithelial lining with pseudostratification. Red arrows, prominent clear cells. (*B*) PicroSirus Red staining shows evidence of marked stromal fibrosis in the corpus of the epididymis. (*C*) p62 accumulation in epithelial cells in Atg5i mice. Clear cells show a stronger accumulation (red arrow).

Together, our data suggest that autophagy deficiency causes degenerative alterations in the epididymis with substantial aggresome accumulation.

### Seminal vesicle and prostate

The SV and prostate are major sources of seminal fluid, which nourish, protect, and help with the transport of sperm. Similar to the epididymis, we observed the accumulation of LC3 and p62 upon Atg5 KD in SV and AP tissues (Supplemental Fig. S1B, Supplemental Fig. S2).

Despite this, H&E staining revealed no obvious histological differences, at least within this time frame (Supplemental Fig. S2). Thus, it is possible that the shrinkage seen in these glands in Atg5i mice is in part attributed to a reduced production of secretory factors (Supplemental Fig. S1A).

To characterise any gene expression changes accompanying autophagy loss in the accessory sex organs, we performed bulk RNA-seq on the SV and AP of 6-week dox-treated experimental and control mice. To further confirm the limited induction of sh-Atg5 in the testis, we included testes in this experiment. As expected, *Atg5* was strongly downregulated in both the AP and SV, but not in the testis, in all experimental samples (Supplemental Fig. S3A). The number and magnitude of the differentially expressed genes (DEGs) caused by Atg5 KD was higher in SV than AP. Interestingly, however, they exhibited a similar trend, implying that autophagy reduction may affect common gene regulatory units across different organs (Fig. 3A). Upregulated DEGs were enriched for inflammatory response and cell cycle genes, among others (Cluster 3&4), suggesting that reduced autophagy causes inflammation and cell proliferation in these tissues, although this was not appreciable in our H&E analysis. Downregulated DEGs also exhibited shared pathways, for instance the unfolded protein response and androgen receptor (AR) pathways.

**Fig. 3.**
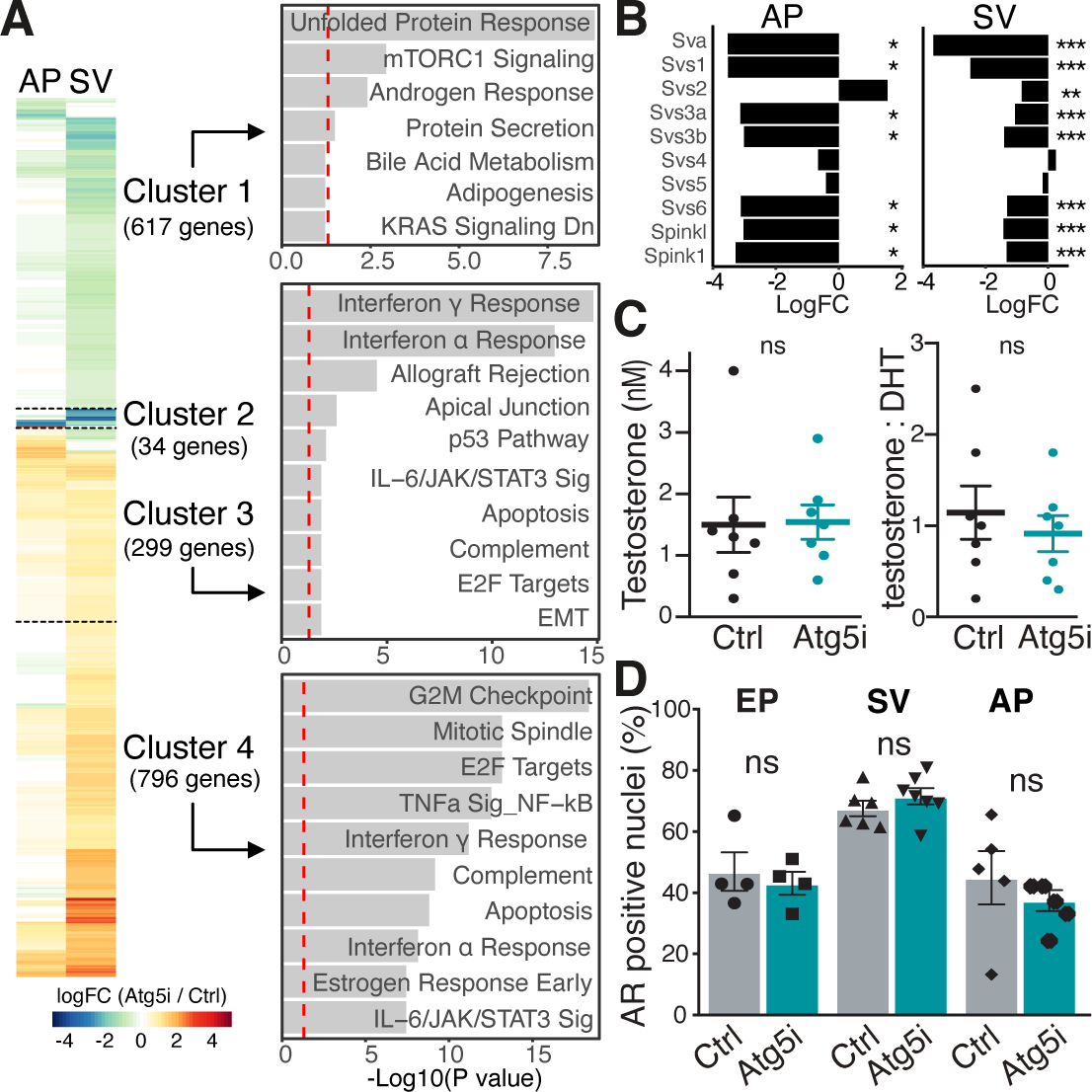
Functional alteration in SV and AP with intact testosterone in Atg5i mice. (*A*) Differentially expressed genes between control and Atg5i (6w on dox) either in seminal vesicles (SV) or anterior prostate (AP). Pathway analysis against MSigDB Hallmark Gene sets at 5% FDR for genes in each of the clusters. (*B*) Altered accessory gland secretions in Atg5i mice. Selected major seminal plasma components are shown, as in (*A*). Bars show log2-fold change, significance FDR after Benjamini-Hochberg correction. (*C*) Tandem mass spectrometry shows no changes in serum testosterone or the testosterone/5α- dihydrotestosterone (DHT) ratio in Atg5i mice. (*D*) Androgen receptor (AR) IHC. Nuclear translocation of AR appears intact in Atg5i mice. EP, epididymis; SV, seminal vesicles; AP, anterior prostate. Values are mean +/- SEM. Student’s t-test.

Notably, several secreted factors, including the *Svs* (seminal vesicle secretions) and *Spink* (serine protease inhibitors of the Kazal type) gene families, were found to be down-regulated (Fig. 3B), thus a reduced fluid production and secretion may account for the decreased organ size observed. These factors are important in forming the copulatory plug in the female mouse, enabling male sperm motility, as well as preventing early acrosomal reaction before the sperm reaches the egg (Kawano et al. 2014; Noda and Ikawa 2019).

It has previously been reported that constitutive loss of autophagy in Atg5^-/-^;NSE-Atg5 mice led to a reduction in circulating testosterone, which may account for the diminished size of the accessory organs and gene expression changes observed in Atg5i mice (Yoshii et al. 2016). To address this, we performed a direct assessment with tandem mass spectrometry on serum isolated from dox-treated Atg5i and control mice. Interestingly, we found total serum testosterone concentrations were comparable to the control mice cohort, with no observable change in the ratio of serum testosterone to dihydrotestosterone (DHT) (Fig. 3C), indicative of normal androgen biosynthesis and activation. This is consistent with the normal size of testes in the dox-treated Atg5i mice (Fig. 1B). Additionally, the nuclear positivity of the androgen receptor (AR) displayed no difference in any tissue analysed (Fig. 3D, Supplemental Fig. S3B). This suggests that, in our Atg5i mice, the loss of autophagy does not result in complete abrogation of the androgen signalling axis, but rather it may partially compromise the signal downstream of the nuclear translocation of AR. Notably, while some *Svs* genes appear AR-responsive (Chen et al. 1987), *Spink1*, which was downregulated in both SV and AP, was reported to be an AR-repressive gene (Tiwari et al. 2020), reinforcing a complex modulation of downstream AR signalling upon Atg5 KD. The apparent discrepancies between these results and the published Atg5^-/-^;NSE-Atg5 mice may reflect the different nature of the systems, wherein the Atg5i model focuses solely on homeostasis and does not incorporate developmental phenotypes.

### Atg5i mice sperm show evidence of oligoasthenozoospermia and reduced fertility

As Atg5i mice display a shrinkage of the accessory sex organs (but not testes) and alterations in seminal fluid production, we wondered whether there would be any impact on sperm and fertility. Sperm were isolated from the epididymis (cauda) for assessment (see methods). Atg5i mice treated with dox for 6 weeks had a significantly reduced sperm count per millilitre compared to controls (Fig. 4A), as ascertained by computer-assisted sperm analysis (CASA), although the percentage of viable sperm cells was similar in both control and Atg5i mice (Fig. 4A). Additionally, using sperm motility grading classifications (Cooper and Yeung 2006), the sperm from control mice had a higher percentage of motile sperm (Grades A and B) than sperm from Atg5i mice (Fig. 4A). Thus, the data suggests that autophagy deficiency in the epididymis may create a sub-optimal environment for sperm maturation. To determine if these alterations influenced fertility, we conducted in vitro fertilization (IVF) efficiency experiments. In agreement, we found there was a significant reduction in fertilisation rates in sperm from dox-treated Atg5i males, as assessed by the presence or absence of two pronuclei at 18-24 hours post IVF (Fig. 4B). Interestingly, the reduced IVF efficiency with Atg5i males was restored in the presence of anti-oxidant reduced glutathione (rGSH) (Ishizuka et al. 2013), further reinforcing the largely intact testis, where spermatogenesis occurs.

**Fig. 4.**
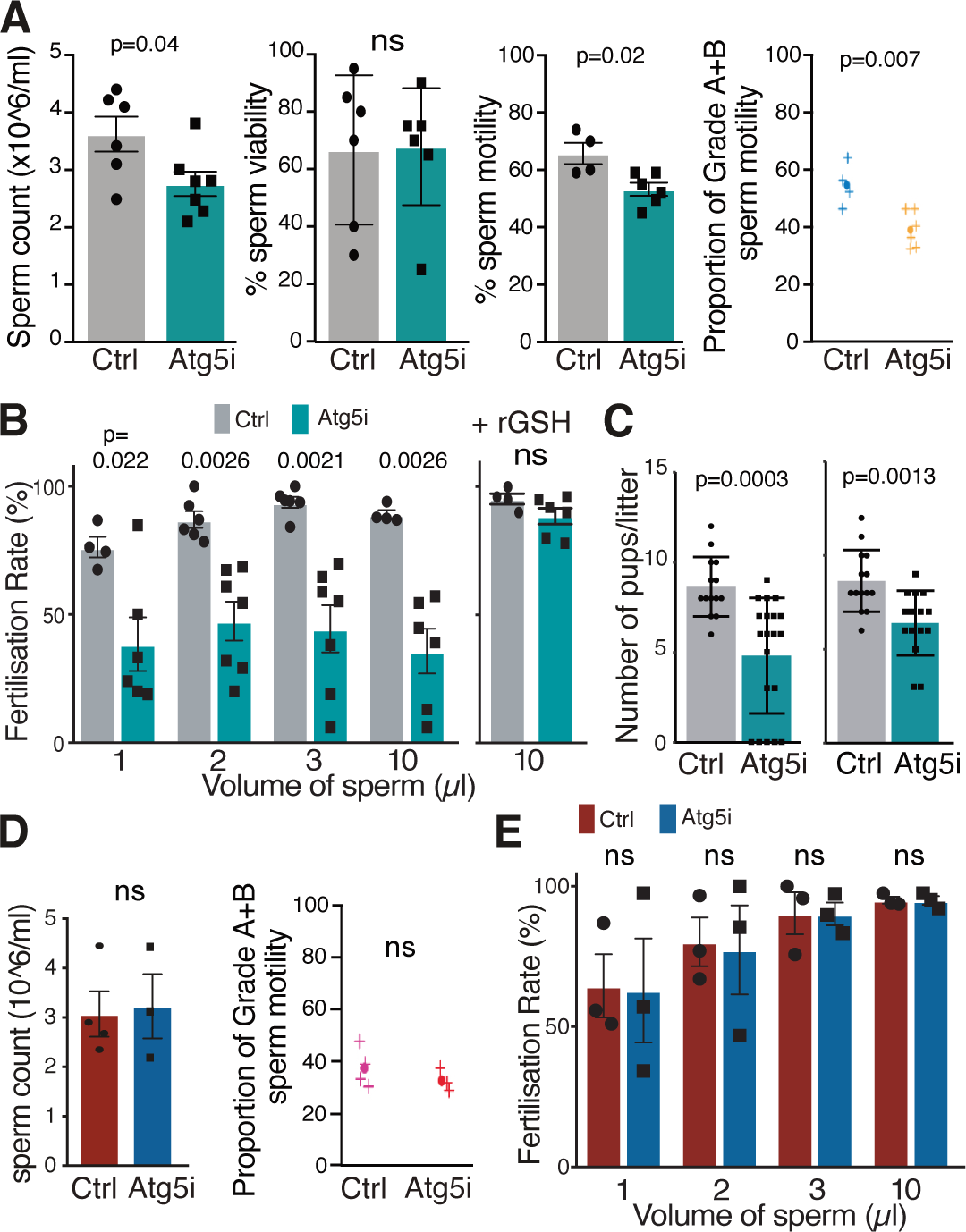
Male subfertility in Atg5i mice. (*A*) Sperm analysis with CASA (see methods): The cauda of the epididymis was dissected for sperm retrieval from control and Atg5i mice (6w on dox). The proportion of moving sperm (Grade A and B) was also plotted. Values are mean +/- SEM. Student’s t-test. (*B*) In vitro fertilisation (IVF) efficiency is based on the formation of two pronuclei using different volumes of sperm with or without anti-oxidant reduced glutathione (rGSH). (*C*) Natural copulation study: 17 fourteen-week-old male mice were bred in ‘trios’ with 34 BL6 female mice (∼10-week-old). All mice were on dox during mating. Once the female showed signs of plug formation, the female was singly-housed and returned to a normal diet. Right, females with no offspring were removed from the analysis. Values are mean +/- SD, p-values Student’s t-test. (*D-E*) Atg5 restoration improves sperm parameters and fertilisation rates in vitro. Eight-week-old mice were given a dox-diet for 6 weeks, followed by a normal diet for additional 6 weeks. Values are mean +/- SEM. Student’s t-test.

We next conducted natural copulation studies using male Atg5i mice (6-weeks on dox) bred to wild-type C57BL/6 females and assessed the number of pups born per litter from a breeding trio. Here, 8/10 male Atg5i mice used in the studies were able to sire pups, but at significantly lower numbers per litter than control mice (Fig. 4C, left), and two male Atg5i mice produced no pups from a trio mating. Atg5i mice experience a decrease in health status upon loss of systemic autophagy (Cassidy et al. 2020), which may deter affected males from mating. However, the reduction in litter size was still significant even after excluding the number of females that did not produce any offspring (Fig. 4C, right). Observations of female copulatory plugs revealed that the females impregnated by control male mice appeared to have thicker and more viscous looking plugs compared to females plugged by male Atg5i mice (Supplemental Fig. S4). This is consistent with the reduced expression in Atg5i mice of genes encoding the Svs proteins, which play a critical role in plug formation in female mice (Fig. 3B). Together, our data suggest that autophagy decline in the post-testicular reproductive organs may cause subfertility, at least in part due to altered sperm maturation and reduced seminal fluid production.

### Autophagy restoration leads to partial restoration of the Atg5i male reproductive phenotype

We next took advantage of the regulable nature of the inducible RNAi system in our mouse model to see if the effects on the accessory sex organs were reversible. Eight-week-old Atg5i and control mice were fed a dox-infused diet for 6-weeks to induce whole-body Atg5 deficiency, and then mice were switched to a standard, dox-free, diet for a further six weeks (Supplemental Fig. S5A). The epididymis, SV, and anterior prostate all showed restoration of Atg5 following a return to a normal, dox-free, diet (Supplemental Fig. S5A). However, the extent to which the Atg5 KD-associated phenotypes recover upon Atg5 restoration varied depending on the tissue type. While the SV and AP from these Atg5-restored mice were found to have almost fully recovered their weight and shape, the epididymis displayed only a partial restoration (Supplemental Fig. S5B).

To determine whether the changes following the restoration of autophagy improved sperm function, we again conducted the sperm analysis and found no differences in the sperm count (Fig. 4D) between Atg5i and control mice. The grade of sperm motility following restoration had a similar proportion of combined Grade A and B sperm (Fig. 4D). Moreover, IVF efficiency following autophagy restoration was also comparable to control mice (Fig. 4E), suggesting that the subfertility associated with temporal inhibition of autophagy is reversible.

Successful reproduction in males mainly involves three essential processes: i) spermatogenesis in the testes, ii) sperm maturation and storage in the epididymis, and iii) semen production in the seminal vesicles and prostate glands. Additionally, there is an overarching hormonal factor known as the pre-testicular step: Gonadotropins released from the pituitary gland stimulate Leydig cells in the testes to produce testosterone, which exerts influence on all these male reproductive organs (Agarwal et al. 2020). Ageing or global autophagy reduction may modulate a complex interrelation between these factors, contributing to the decline of sperm parameters; however, it is challenging to dissect the individual steps to determine the cause of male subfertility. The Atg5i mice provide a unique opportunity in this regard, likely due to the BTB: in male Atg5i mice on dox (e.g., for 6w), whereas the gross appearance of testes was unaffected, the accessory sex organs exhibited a dramatic alteration, reminiscent of ‘castration or androgen deprivation’. Importantly, Atg5 mice also showed intact testosterone levels, thus allowing us to study the specific role of autophagy in post-testicular organs. The precise reason for the atrophic appearance of these organs remains unclear. As suggested by our RNA-seq data, it is likely that a dramatic reduction in seminal fluid production occurs. While upstream AR signalling appears intact, it is also possible that modulation of downstream AR signalling plays a role. Notably, Leydig cells, the major testosterone-producing cells, sit in the interstitial space outside of the BTB, and thus can express sh-Atg5 upon dox-treatment in this model. It was shown that depletion of *Atg5* or *Atg7* in Leydig and other steroidogenic cells in mice leads to a defect in cholesterol uptake and reduced testosterone production (Wang et al. 2021; Gao et al. 2018). Thus, we do not entirely rule out the involvement of testicular factors in our model. It is possible that the hypomorphic nature of RNAi may contribute to the uniqueness of our model.

## Author contributions

A.J., S.P. and M.N conceived the project. A.J., L.D.C., and A.R.J.Y. performed in vivo experiments. A.T. and W.A. undertook steroid analysis. A.Y.W. conducted pathology. C.L.R.B. analysed data and supervised the study. G.L. performed IVF. M.L.B. and O.D. performed sperm analysis. A.S.L.C. performed RNA-seq analysis. A.J., L.D.C., A.R.J.Y., and M.N. wrote the manuscript, and all authors read and edited the manuscript.

## Materials and Methods

### Mice

The generation and initial characterisation of the Atg5i transgenic mice was previously described (Cassidy et al. 2018). Mice were maintained on a mixed C57BL/6 × 129 background with littermate controls used in all experiments.

### Immunohistochemistry (IHC)

IHC was performed as previously (Cassidy et al. 2018). Tissue samples were fixed in 10% neutral buffered formalin for 24 hours, transferred to 70% ethanol, machine processed (Leica Asp300 Tissue Processor; Leica, Wetzlar, Germany), and embedded in paraffin. Formalin-fixed paraffin-embedded (FFPE) samples were de-waxed and rehydrated. Heat-induced antigen retrieval was performed in a pressure cooker for 5 min at 120 °C in either citrate buffer (10 mM sodium citrate, 0.05% Tween 20, pH 6) for p62 (Enzo, BML-PW9860, 1;750), LC3 (Abcam, ab128025, 1:1000 for epididymis or 1:2000 for SV) and v-ATPase (Merck, HPA028701, 1:1000) or Tris-EDTA buffer (10mM Tris Base, 1mM EDTA Solution, 0.05% Tween 20, pH 9) for AR (Santa Cruz, sc-816, 1:500) and tGFP (Thermo Fisher, PA5-22688, 1:1000).

### Picrosirius red staining

To stain for collagen, FFPE slides were de-waxed and rehydrated before being stained with Weigers Haematoxylin for 8 minutes (CellPath RBA-4201-00A), washed in running water for 10 minutes, immersed in picrosirius red solution 0.1% [w:v] Direct Red 80 (Sigma 365548) in a saturated picric acid solution (Sigma P6744) for 1 hour and washed in acidified water (0.5% acetic acid) twice. Slides were then dehydrated and cover-slipped.

### Western

Western was performed as described before (Cassidy et al. 2020). Tissue samples were homogenised with the Precellys 24 tissue homogeniser in Laemmli buffer. Following SDS-PAGE electrophoresis, the proteins were transferred to a PVDF membrane (Immobilon, Millipore), which was subsequently blocked for 1 h at room temperature (5% Marvel milk (Premier foods, Dublin, Ireland) solution in TBS-Tween 0.1%) before incubating with primary antibody at 4 °C overnight. The following primary antibodies were used: anti-β-Actin (Sigma, A5441, 1:10,000); anti-ATG5 (Abcam, ab108327, 1:1000); and anti-LC3 (Abcam, ab192890, 1:1000).

### RNA-seq

RNA was extracted using the Qiagen RNeasy plus kit (Cat #74136) according to the manufacturer’s instructions and quality checked using a Bioanalyser Eukaryote Total RNA Nano Series II chip628 (Agilent, Cat # 5067-1511). Libraries were prepared using the TruSeq Stranded mRNA Library Prep Kit (Illumina Cat # 20020594) according to manufacturer’s instructions and sequenced using the HiSeq-2500 platform (Illumina).

Reads were aligned to the mouse genome, version GRCm38, using STAR version 2.7.6a (Dobin et al. 2013) and per-gene read counting was performed using the featureCounts function of the subread version 1.5.3 (Liao et al. 2013). Differential expression analysis was performed with edgeR (McCarthy et al. 2012), comparing each of the Atg5i samples with their control equivalent from the same tissue. DE genes were identified using edgeR’s glmTreat function (McCarthy et al. 2012) using a fold change of 1.2 in either direction and an FDR cutoff of 0.05.

Overlap-based pathway and gene ontology enrichment was performed using the web-based Enrichr platform (Chen et al. 2013; Kuleshov et al. 2016). Differentially expressed genes were tested against genesets curated as part of the Molecular Signatures Database (MSigDB) (Liberzon et al. 2011, 2015).

### Sperm analysis

Sperm were isolated from the cauda region of the epididymis. Sperm count, motility and morphology were assessed using a Sperminator Computer-Aided Sperm Analysis (CASA) system linked to a microscope with a heated stage. Assessment of sperm concentration and motility was performed at 36°C using a heated stage. An aliquot of the sperm sample was added to a CellVision slide (CellVision Technologies, Netherlands) using a positive displacement pipette. The CellVision slide was placed onto the heated stage and analysed after a period of 2 minutes to allow the spermatozoa to settle.

The CASA system was used to assess the sperm concentration and sperm motility. It was set up according to the manufacturer’s instructions and the sperm was visualised and tracked. The concentration and motility were divided into four categories:

**A:** excellent or rapid progression, sperm moving actively at more than half a tail length per second.

**B:** sluggish progression, sperm moving actively but slower than half a tail length per second. **C:** non-progressive motility, all other categories of motility with an absence of progression including swimming in circles or weak flagellar force.

**D:** not motile or immobile, static sperm with no movement.

For calculating progressive motility, it is recommended for greater accuracy and consistency to combine Grade A and B (Cooper and Yeung 2006; Cooper et al. 2010).

For the morphology assessment, sperm was fixed onto glass sides. Sperm was then viewed with a Leica TCS SP5 microscope at x40 magnification. Different sperm forms were assessed and classified as per Bourn Hall in-house sperm morphological assessment protocol as described below.

**Normal forms:** a smooth, regularly contoured and generally hooked shaped head. The acrosome is well defined in mice. The post acrosomal region contains no vacuoles. The mid- piece is slender and the tail is of a uniform thickness, thinner than the mid-piece and with no sharp angles.

**Head defects:** The hook-shape has been lost and the head appears flat or round. It may also appear vacuolated or have a double head.

**Neck/midpiece defects:** asymmetrical insertion of the mid-piece into the head, thick or irregular, sharp bent, abnormally thin (subjective), or in any of these combinations.

**Tail defects:** short, multiple, broken, smooth hairpin bends, sharply angulated bends, irregular width, coiled, or any of these combinations.

Morphological assessments were performed by averaging the number of normal and abnormal forms from at least two random sections on the slide. The average percentage was then reported.

### In vitro fertilisation (IVF)

10-14 weeks old female C57BL/6J mice were intraperitoneally (IP) injected with 7.5 IU Pregnant Mare’s Serum Gonadotropin (PMSG), and then re-injected IP with 7.5 IU human chorionic gonadotropin (hCG) 48 hours later. The oocytes were subsequently harvested from the super-ovulated C57BL/6J mice approximately 14 to 16 hours post hCG injection.

490 μL, 495 μL, 498 μL and 499 μL of tubule fluid (TF) medium with or without reduced glutathione (rGSH) (Sigma-Aldrich, G-4251) at final1.25mM were added to individual wells of a 4-well plate. Sperm released from the epididymis was placed in 500 μL cryoprotectant agents (CPA) medium. This was then incubated at 37°C, 5% CO2 for 10 minutes. 10 μL, 5 μL, 2 μL and 1 μL CPA medium containing sperm was then added to the already prepared 4 wells of the 4-well plate containing the TF medium to give a total volume of 500 μL in each well. The 4-well plate was then placed in an incubator and allowed to equilibrate for at least 45 minutes prior to oocyte collection.

Next, oviducts from 5 female mice were dissected in a small dish (35 mm) containing 10 mL TF solution on a heated plate (37°C). The cumulus masses from the swollen ampulla were released into TF medium and 4 to 5 cumulus masses were transferred into each IVF well of the 4 well-plate. The 4 well-plate was placed in an incubator at 37°C, 5% CO2.

After 3 to 4 hours of culturing in the incubator, the 4-well plate was removed and working under a microscope, the oocytes were rinsed three times to thoroughly remove cell debris, degenerating oocytes and dead sperm. KSOM medium was added to the 4-well plate, which was then incubated at 37°C, 5% CO2, overnight. Finally, the number of polar bodies corresponding to the fertilisation rate was recorded.

### Steroid analysis

Serum testosterone and 5α-dihydrotestosterone were measured by liquid chromatography-tandem mass spectrometry (LC-MS/MS) using a Waters Xevo mass spectrometer as previously described (Schiffer et al. 2022; O’Reilly et al. 2014). In brief, steroids were extracted from 200 μL of mouse serum. 20 μL of an internal standard solution (testosterone-d3 and DHT-d3) was added, and steroids were extracted via liquid/liquid extraction using 2mL of methyl-tert butyl ether. The organic layer was removed and evaporated to dryness under nitrogen at 55 °C Samples were reconstituted in 125 μL of 50/50 methanol/water for mass spectrometry analysis. Steroids were quantified relative to a calibration series ranging in concentration from 0.25 to 500 ng/mL (with inclusion of a blank).

### Data availability

RNA-seq data have been deposited in GEO with accession number GSE235690.

## Acknowledgements

This work was supported by a Cancer Research UK Cambridge Institute core (no. C9545/A29580). We are grateful to the following CRUK Cambridge Institute core facilities for advice and assistance: Histopathology, light microscopy, genomics, bioinformatics, genome editing, and the BRU. M.N. is also supported by BBSRC (BB/S013466/1, BB/T013486/1) and Diabetes UK via BIRAX and the British Council (65BX18MNIB). A.J. was supported by a Cambridge Cancer Centre Clinical PhD Research Fellowship. LDM was supported by BBSRC (BB/T013486/1). AYW is supported by the Cancer Research UK Cambridge Centre (C9685/A25177) and NIHR Cambridge Biomedical Research Centre (BRC-1215-20014). W.A. is supported by the Wellcome Trust (Investigator Award 209492/Z/17/Z).

---

**Supplemental Fig. S1. ‘Castration-like’ phenotype in Atg5i mice.** (*A*) Representative picture of male reproductive organs in indicated mice (6w on dox). SV; Seminal Vesicle, AP; Anterior Prostate. (*B*) Western blotting for indicated proteins. (*C*) Representative IHC images for LC3 in the epididymis. (*D*) Representative IHC images for p62 and v-ATPase, a marker of clear cells in the epididymis, caput. Serial sections were used (Atg5i). Square regions are magnified (lower left). Red arrows indicate clear cells.

**Supplemental Fig. S2. Histological analysis of seminal vesicle and anterior prostate in Atg5i mice.** Representative images of H&E and indicated IHC in control and Atg5 mice (6w on dox).

**Supplemental Fig. S3. Gene regulation in seminal vesicles and anterior prostates in Atg5i mice.** (*A*) Volcano plots of differentially expressed genes between control and Atg5i mice (6w on dox) in the indicated tissues. (*B*) Representative images of IHC for androgen receptor (AR) in indicated samples.

**Supplemental Fig. S4. Natural copulation in male Atg5i and wild-type female mice.** Representative images show that females impregnated by Control male mice have thicker and more viscous-looking plugs as compared to females plugged by Atg5i mice.

**Supplemental Fig. S5. Partial reversibility of accessory sex organ phenotype in male Atg5i mice.** (*A*) Eight-week-old mice were given a 6-week dox diet. At 14 weeks old, they were returned to a normal diet for 6 weeks. Then Western blotting was performed for indicated proteins. (*B*) After the autophagy recovery, tissue morphology and weight were assessed. Values are mean +/- SEM. Student’s t-test.

