## Supplemental Figs 1-5 for "Autophagy maintains the homeostatic environment in the male reproductive accessory organs playing a key role in fertility"

### Supple_Fig. 1 _ Jaulim et al.

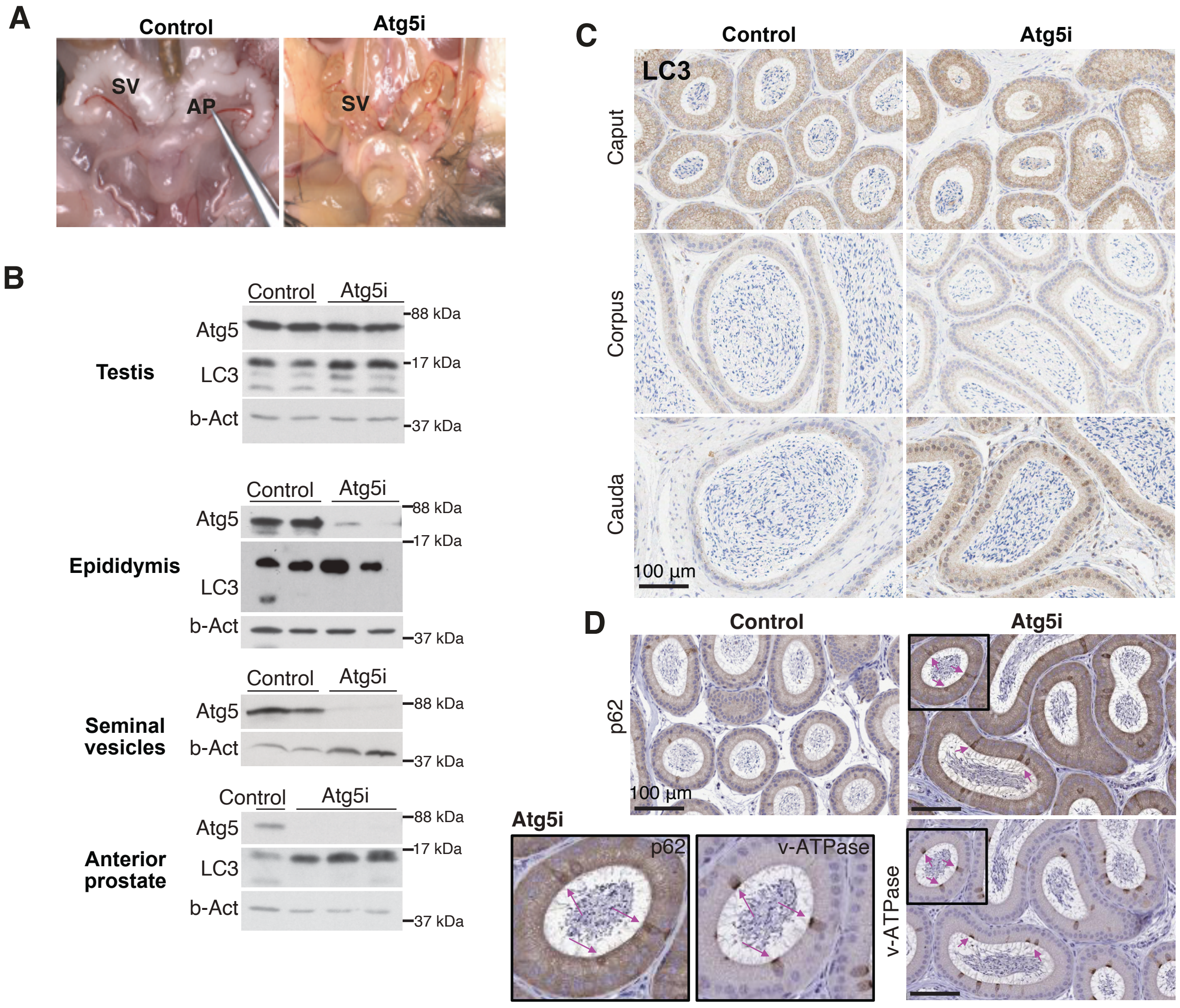


**Supplemental Fig. S1. 'Castration-like' phenotype in Atg5i mice.** (*A*) Representative picture of male reproductive organs in indicated mice (6w on dox). SV; Seminal Vesicle, AP; Anterior Prostate. (*B*) Western blotting for indicated proteins. (*C*) Representative IHC images for LC3in the epididymis. (*D*) Representative IHC images for p62 and v-ATPase, a marker of clear cells in the epididymis, caput. Serial sections were used. Square regions are magnified (lower left). Red arrows indicate clear cells.

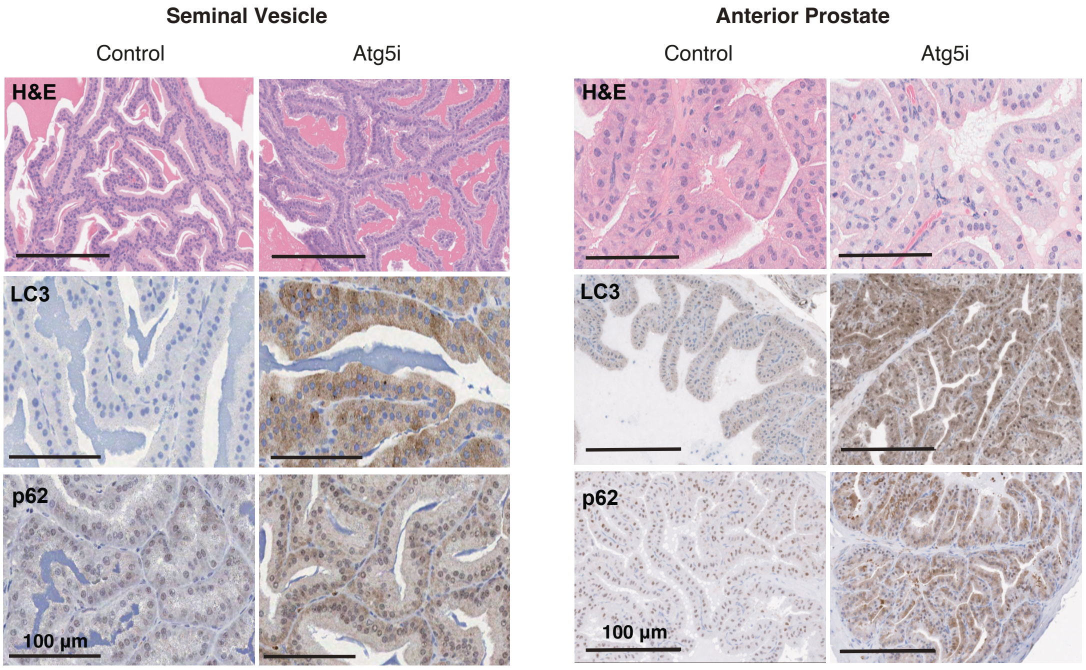


**Supplemental Fig. S2. Histological analysis of seminal vesicle and anterior prostate in Atg5i mice.** Representative images of H&E and indicated IHC in control and Atg5 mice (6w on dox).

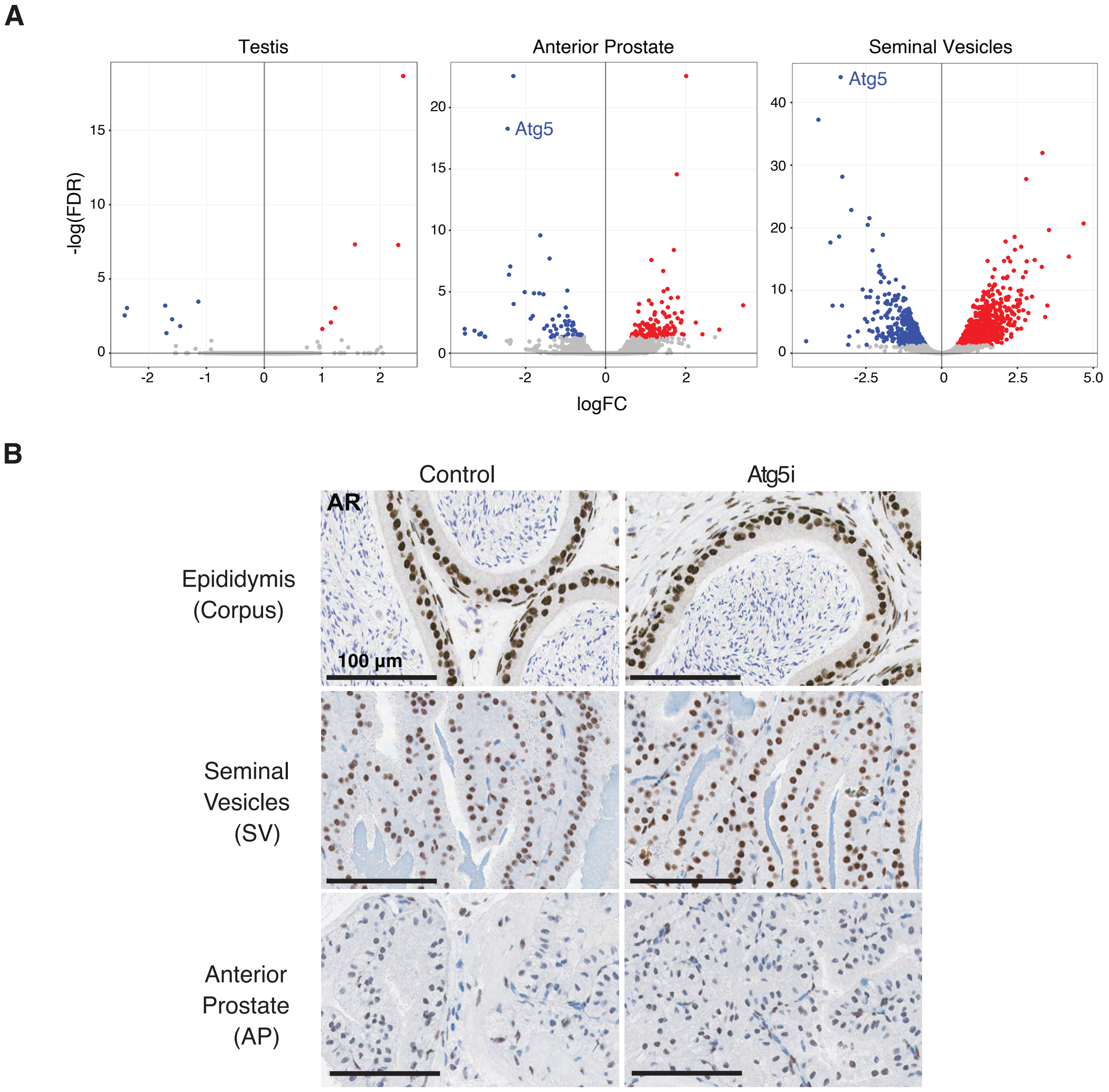


**Supplemental Fig. S3. Gene regulation in seminal vesicles and anterior prostates in Atg5i mice.** (*A*) Volcano plots of differentially expressed genes between control and Atg5i mice (6w on dox) in the indicated tissues. (*B*) Representative images of IHC for androgen receptor (AR) in indicated samples.

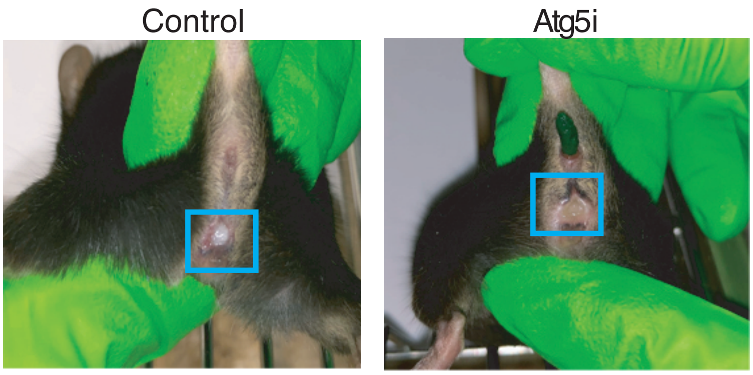


**Supplemental Fig. S4. Natural copulation in male Atg5i and wild-type female mice.** Representative images show that females impregnated by Control male mice have thicker and more viscous-looking plugs as compared to females plugged by Atg5i mice.

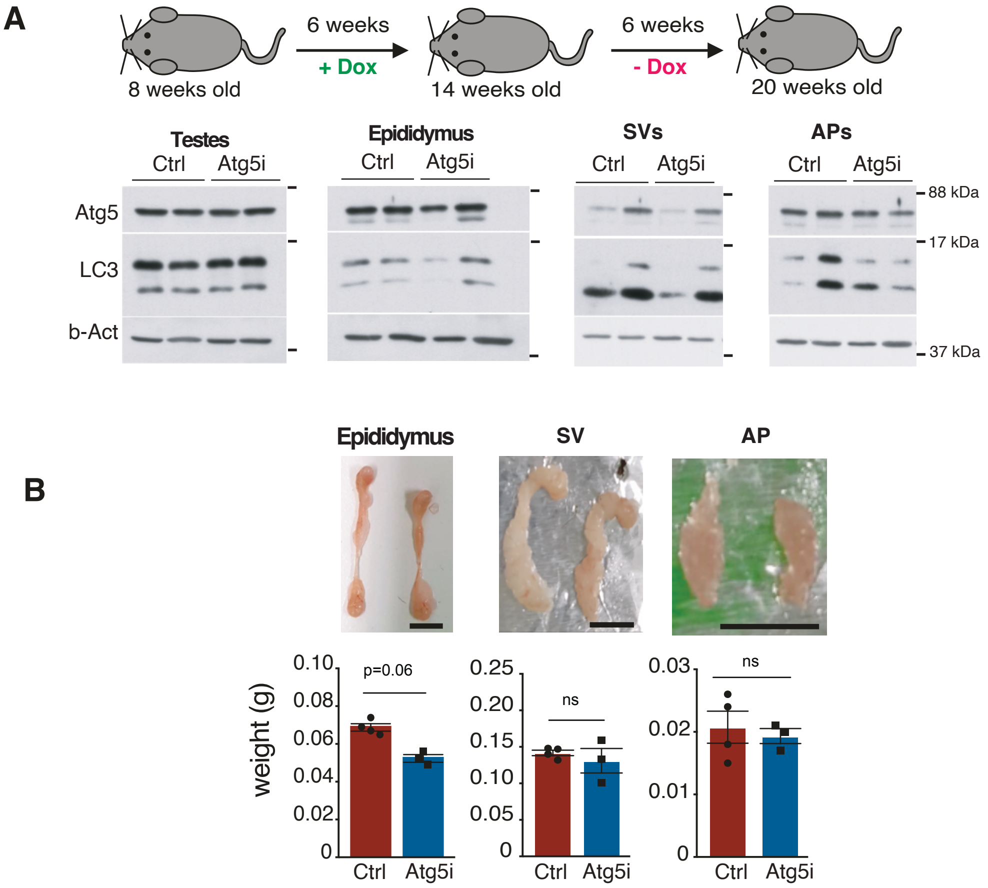


**Supplemental Fig. S5. Partial reversibility of accessory sex organ phenotype in male Atg5i mice.** (*A*) Eight-week-old mice were given a 6-week dox diet. At 14 weeks old, they were returned to a normal diet for 6 weeks. Then Western blotting was performed for indicated proteins. (*B*) After the autophagy recovery, tissue morphology and weight were assessed. Values are mean +/- SEM. Student's t-test.
